# Genetic and environmental determinants of variation in the plasma lipidome of older Australian twins

**DOI:** 10.1101/2020.05.05.075606

**Authors:** Matthew W.K. Wong, Anbupalam Thalamuthu, Nady Braidy, Karen A. Mather, Yue Liu, Liliana Ciobanu, Bernhardt T. Baune, Nicola J. Armstrong, John Kwok, Peter R. Schofield, Margaret J. Wright, David Ames, Russell Pickford, Teresa Lee, Anne Poljak, Perminder S. Sachdev

**Author notes:** denotes equal last author (Anne Poljak, Perminder S. Sachdev). Corresponding author: Prof. Perminder S. Sachdev.

## Abstract

The critical role of blood lipids in a broad range of health and disease states is well recognised, while an understanding of the complex genetic regulation of lipid homeostasis is emerging. Traditional blood lipids (LDL-C, HDL-C and triglycerides) are known to be substantially regulated by genetic variation. Less well explored is the interplay of genetics and environment within the broader blood lipidome. Here we use the twin model to examine heritability of the plasma lipidome among healthy older aged twins and explore gene expression and epigenetic (DNA methylation) associations of these lipids. Heritability of 209 plasma lipids quantified by liquid chromatography coupled mass spectrometry (LC-MS) was assessed in 75 monozygotic and 55 dizygotic twin pairs enrolled in the Older Australian Twins Study (OATS), aged 69-93 years. Only 27/209 lipids (13.3%) were significantly heritable under the classical ACE twin model (*h*^2^ = 0.28-0.59). Ceramides (Cer) and triglycerides (TG) were most heritable, while sphingomyelins (SM) and most phospholipids, especially lysophospholipids, were not significantly heritable. Lipid levels correlated with 3731 transcripts. Relative to non-significantly heritable TGs, heritable TGs had a greater number of associations with gene transcripts, which were not directly associated with lipid metabolism, but with immune function, signalling and transcriptional regulation. Genome-wide average DNA methylation (GWAM) levels accounted for a proportion of variability in some non-heritable lipids, especially lysophosphatidylcholine (LPC). We found a complex interplay of genetic and environmental influences on the ageing plasma lipidome, with most of the variation controlled by unique environmental influences.

## Introduction

As the field of lipidomics has grown, hundreds to thousands of complex lipids have been characterised ^1; 2^, with many linked to health and disease states, such as metabolic syndrome ^3^, cardiovascular disease ^4; 5^, obesity ^6; 7^, and dementia ^8–11^. Both genetic and environmental factors influence these biological phenotypes. Identifying the contributions of these factors can help elucidate the importance of genes for a particular trait, as well as providing insight into the environmental influences. This information might enable the design of personalised medical treatments for lipid-related disease states.

While there are substantial data to suggest that levels of traditional lipids and lipoproteins such as high density lipoprotein (HDL) cholesterol, low density lipoprotein (LDL), cholesterol and triglyceride levels are heritable ^12; 13^, few studies have focused on the genetic and environmental influences on the plasma levels of individual lipid species and lipid classes beyond these traditional lipid measures. Additionally, lipids vary within and between individuals ^14–16^ based on variables such as age ^17–19^, sex ^17; 19^, body mass index (BMI) ^19; 20^, lipid-lowering medication ^21^ and genetic background ^12; 22^, demonstrating a wide degree of complexity involved in the regulation of lipid metabolism. It would therefore be informative to understand the extent to which variation in specific plasma lipids is determined by genetic and environmental influences. We hypothesise that as circulating lipids are produced downstream of genomic, transcriptomic and proteomic regulatory processes, that there will be strong environmental influences on lipid variance.

Previous genome-wide association study (GWAS) data implicate many genetic loci associated with traditional lipid levels. For example, the genes encoding lipoprotein lipase, hepatic lipase and cholesteryl ester transfer protein (*LPL, LIPC* and *CETP* respectively) have been associated with HDL, and genes encoding cadherin EGF LAG seven-pass G-type receptor 2, apolipoprotein B and translocase of outer mitochondrial membrane 40 (*CELSR2, APOB* and *TOMM40* respectively) have been associated with LDL ^23^. Apolipoprotein E (*APOE*) variants have been established as a strong risk factor for cardiovascular disease and Alzheimer’s disease ^22; 24^ and are associated with altered LDL-C levels. One large exome wide screening study with over 300,000 individuals identified 444 variants at 250 loci to be associated with one or more of plasma LDL, HDL, total cholesterol and triglyceride levels ^25^. Collectively, data from 70 independent GWAS with sample sizes ranging from ten thousand to several hundred thousand participants have identified associations of traditional lipid levels with 500 single nucleotide polymorphism (SNPs) in 167 loci that explain up to 40% of individual variance in these traditional lipid measures ^26^. This number suggests that LDL, HDL, total cholesterol and triglyceride levels undergo a substantial degree of genetic regulation, but also highlights that much of the lipid variance is still unaccounted for, possibly related to rare variants or environmental factors ^26; 27^.

One of the most powerful tools for analysis of gene versus environment effects on phenotypic traits is the classical twin design, which estimates the relative contribution of heritable additive genetic effects (A) and shared (C) and unique environmental (E) influences on a given trait by comparing correlations within monozygotic and dizygotic twin pairs ^28^. One major strength of this design compared to family studies is that twins are matched by age and common environment, reducing cross-generation differences. Genetic and environmental variances can be computed with relatively high power using a modest sample size. It is expected that since monozygotic twins share 100% of segregating genetic variation, while dizygotic twins share 50%. It is also assumed that twins are raised in the same environment, thus any additional differences between monozygotic twins would be attributable to unique environmental (E) effects. Further, any differences in intraclass correlations between monozygotic and dizygotic twins could be estimated as due to additive polygenic effects (A).

We applied the classic twin design to estimate heritability using 75 pairs of MZ twins and 55 pairs of DZ twins from the Older Australian Twin Study (OATS) ^29; 30^, aged between 69-93 years. Since many proteins are known to regulate lipid metabolism, it is expected that some lipids may show substantial heritability, as reported in previous studies ^31; 32^. Further, we hypothesised that some of the variance in lipids that do not have significant heritability might be controlled by gene sequence - independent mechanisms, such as genome-wide average DNA methylation (GWAM) levels. Our study is the first to examine heritability of the broad plasma lipidome among healthy older – aged twins and explore putative genetic, transcriptomic and epigenetic associations of these lipids.

## Materials and Methods

### Cohorts

The study sample comprised participants aged between 69-93 years enrolled in the Older Australian Twin Study (OATS), established in 2007. The study recruited participants from three states in eastern Australia (QLD, NSW and VIC). The OATS collection included; patient data, including blood chemistry, MRI, neuropsychiatric assessment/cognitive tests, and medical exams performed over several visits (waves), each taken at an interval of 16-18 months, with the first visit denoted as “Wave 1”, second visit denoted as “Wave 2” and so on. From OATS, we selected *n*=330 participants who had available plasma from Wave 3; plasma from this wave collected within a period of up to 3 years apart. Of these, 260 participants were eligible for heritability analyses, including 150 monozygotic twins (75 pairs in total; 25 male, 50 female), and 110 dizygotic twins (55 pairs in total; 31 males, and 79 females). The study protocol for OATS has been previously published ^29; 30; 33^. Participants who had significant neuropsychiatric disorders, cancer, or life threatening illness were excluded from this study.

### Ethics Approval

OATS was approved by the Ethics Committees of the University of New South Wales and the South Eastern Sydney Local Health District (ethics approval HC17414). All work involving human participants was performed in accordance with the principles of the Declaration of Helsinki of the World Medical Association. Informed consent was obtained from all participants and/or guardians.

### Plasma collection, handling and storage

Blood collection, processing and storage were performed under strict conditions to minimize pre-analytical variability ^11^. Fasting EDTA plasma was separated from whole blood within 2-4 hours of venepuncture and immediately stored at −80°C prior to bio-banking. Samples then underwent a single freeze thaw cycle for the purpose of creating aliquots, which minimizes subsequent freeze thaw cycles for specific experiments. EDTA plasma was chosen as the anticoagulant since it chelates divalent metals, thereby protecting plasma constituents from oxidation, which is particularly important for lipids. Thereafter, lipid extractions were performed within 15 minutes of freeze thawing and extracts stored at −80°C and analysed within two months of extraction.

### Targeted assays of plasma lipids

Plasma total cholesterol, LDL-C, HDL-C and TG were measured by enzymatic assay at SEALS pathology (Prince of Wales Hospital) as previously described ^34^, using a Beckman LX20 Analyzer with a timed-endpoint method (Fullerton, CA). LDL-C was estimated using the Friedewald equation (LDL-C =total cholesterol - HDL-C - triglycerides/2.2).

### *APOE* genotyping

DNA was extracted from samples using established procedures ^35^. Genotyping of two *APOE* single nucleotide polymorphisms (SNPs rs7412, rs429358) was performed using Taqman genotyping assays (Applied Biosystems Inc., Foster City, CA) to determine the *APOE* haplotype, which has three alleles (ε2, ε3, ε4).

### Lipid Extraction from plasma: Single phase 1-butanol/methanol

Lipid internal standards (SPLASH® Lipidomix® Mass Spec Standard) were purchased from Avanti (Alabaster, Alabama, United States) and diluted ten-fold in 1-butanol/methanol (1:1 v/v). Plasma extraction was performed in accordance with a single phase extraction as previously described ^36; 37^. Briefly, we added 10 μL of 1:10 diluted SPLASH internal lipid standards mixture to 10 μL plasma in Eppendorf 0.5 mL tubes. 100 μL of 1-butanol/methanol (1:1 v/v) containing 5 mM ammonium formate was then added to the sample. Afterwards, samples were vortexed for 10 seconds, then sonicated for one hour. Tubes were centrifuged at 13,000 g for 10 minutes. The supernatant was then removed via a 200 μl gel-tipped pipette into a fresh Eppendorf tube. A further 100μl of 1-butanol/methanol (1:1 v/v) was added to the pellet to re- extract any remaining lipids. The combined supernatant was dried by vacuum centrifugation and resuspended in 100 μl of 1-butanol/methanol (1:1 v/v) containing 5 mM ammonium formate and transferred into 300 μl Chromacol autosampler vials containing a glass insert. Samples were stored at −80° C prior to LC-MS analysis. The robustness and reproducibility of this extraction method has been previously demonstrated ^37^ in our laboratory, with variation in human plasma ranges of measurement between individuals across age, sex ^19^ and by *APOE* genotype ^38^ reported.

### Liquid Chromatography/ Mass spectrometry

Lipid analysis was performed by LC ESI-MS/MS using a Thermo QExactive Plus Orbitrap mass spectrometer (Bremen, Germany) in two experimental batches separated by a month. A Waters ACQUITY UPLC CSHTM C18 1.7um, 2.1×100mm column was used for liquid chromatography at a flow rate of 260 μL/min, using the following gradient condition: 32% solvent B to 100% over 25 min, a return to 32% B and finally 32% B equilibration for 5 min prior to the next injection. Solvents A and B consisted of acetonitrile:MilliQ water (6:4 v/v) and isopropanol:acetonitrile (9:1 v/v) respectively, both containing 10 mM ammonium formate and 0.1% formic acid. Product ion scanning was performed in positive ion mode. Sampling order was randomised prior to analysis.

### Alignment and peak detection/analysis

The raw data was aligned, chromatographic peaks selected, specific lipids identified and their peak areas integrated using Lipidsearch software v4.2.2 (Thermo Fischer Scientific, Waltham MA). Owing to the large number of RAW files being processed, the alignment step was performed in four separate batches, with a maximum of 100 samples aligned at any one time, and the data collated and exported to an Excel spreadsheet for manual processing and statistical analysis. Only lipids that were present in all four alignment batches were included in our analysis. The raw abundances (peak areas) were normalised by dividing each peak area by the raw abundance of the corresponding internal standard for that lipid class e.g. all phosphatidylcholines were normalised using 15:0-18:1(d7) PC. The intra-assay coefficient of variation (CV) was calculated by dividing the standard deviation of the normalised abundances by the mean across lipid species. Lipid ion identifications were filtered using the LipidSearch parameters rej=0 and average peak quality>0.75. Furthermore, identifications with CV<0.4 from repeated injections of quality control plasma samples were included (see S1 supporting methods). Where duplicate identifications were found (i.e. lipid IDs with identical m/z and annotations, and similar retention times), the lipid ID with the lowest CV%, and highest peak quality score was used. When necessary, the average m-score (match score, based on number of matches with product ion peaks in the spectrum [20]) was also used to differentiate closely related lipid species, with the lipid having the highest m-score selected. All other duplicates were excluded from analysis. Lipid groupsums were produced by adding lipids within a defined class/subclass together, such as total monounsaturated triglycerides (TG), total ceramides (Cer) etc.

### Microarray Gene Expression

Fasting blood samples for gene expression analyses were collected. The methods for gene expression data collection analyses have previously been described ^39^. Briefly, PAXgene Blood RNA System (PreAnalytiX, QIAGEN) was used to extract total RNA from whole blood collected in PAXgene tubes following overnight fasting. RNA samples with RNA integrity number (RIN) ≥6 as measured by the Agilent Technologies 2100 Bioanalyzer were used in subsequent analyses ^40^. Assays for gene expression were performed using the Illumina Whole-Genome Gene Expression Direct Hybridization Assay System HumanHT-12 v4 (Illumina Inc., San Diego, CA, USA) in accordance with standard manufacturer protocols. Quality control (QC) and pre-processing of raw gene expression intensity values extracted from GenomeStudio (Illumina) were performed using the R Bioconductor package limma ^41^. Background correction and quantile normalisation was done using the neqc function. Expressed probes with detection p-value <=0.05 were retained for analysis. After pre-processing and filtering, 308 samples and 36,053 transcripts were available for gene expression analysis. After overlapping with the lipids data 290 samples were available for lipids – gene expression analysis. Gene abbreviations used in the text are based on Gene Ontology nomenclature.

### DNA Methylation

Genome-wide DNA methylation data for 113 monozygotic twin pairs was generated using an established genomics provider using peripheral blood DNA collected at baseline ^42^. Randomisation of co-twins across the arrays was performed within experiments. DNA methylation status was assessed using the Illumina Infinium HumanMethylation450 BeadChip (Illumina Inc., San Diego, CA, USA). Background correction was applied to raw intensity data and the R *minfi* package was used to generate methylation beta values (ranging from 0-1) ^43^. Quantile normalisation was used. We excluded sex chromosome probes, probes containing SNPs, cross-reactive probes as well as probes not detected in all samples from analysis ^44^. Following these quality control (QC) procedures, 420,982 out of 485,512 probes remained. White blood cell composition was estimated using a previously described method ^45^, implemented in *minfi*. After filtering methylation outliers using the preprocessQantile function of the minfi package with default parameters, out of the 217 samples with methylation data, 135 overlapped with lipids data. Genome wide Average Methylation (GWAM) for each sample across all the probe level beta values were calculated.

### Data Analysis

#### Data Transformations

Since different sets of covariates are used to adjust for the lipid levels, gene expression and methylation, we have first obtained residuals after adjusting for standard confounders in order to obtain lipid and gene expression profiles independent of cohort characteristics. Residuals for lipids were obtained after adjusting for age, sex, education, BMI, lipid lowering medication, smoking status, experimental batch and *APOE* ε4 carrier status, which were then inverse normal transformed using the R package RNOmni ^46^. This transformation eliminated experimental batch separation effects (Figure S2). Residuals for gene expression were obtained after adjusting for age, sex, experimental batch, RIN, blood cell counts (eosinophils, lymphocytes, basophils and neutrophils - obtained using standard laboratory procedures by Prince of Wales SEALS Pathology). Residuals for methylation beta values were obtained after adjusting for age, sex, BMI and estimated white blood cell counts (CD8T, CD4T, NK, B-cell, monocytes, and granulocytes). Residuals were used for all the analyses presented here.

#### Heritability Estimation

Heritability was estimated using SEM. Under the SEM the phenotypic covariance between the twin pairs is modelled as a function of additive genetic (A), shared environmental (C) and unique environmental (E) components. In the narrow sense heritability is defined as the ratio of additive genetic variance [Var(A)] to the total phenotypic variance [Var(A)+Var(C)+Var(E)]. The model containing these three parameters (A, C and E) is known as the ACE model. For model parsimony and test concerning the variance parameters, models with only A and E components, known as AE model, and the models with CE and E components would be fit and compared with the full ACE model ^47^. Genetic and environmental correlations were estimated using the bivariate Cholesky model. Heritability, genetic correlations and environmental correlations under the twin SEM were estimated using the R OpenMx (2.12.1.) package ^48^.

#### Association Tests

Test of association of lipids with probe level gene expression were performed using the linear mixed model and the lme function in R package nlme ^49^. Gene expression and lipid residuals (adjusted for age, sex, education, BMI, lipid lowering medication, smoking status, experimental batch and *APOE* ε4 carrier status) were used as independent and dependent variables respectively in these models. A p-value threshold of 1.39×10^−6^ (0.05/35971, obtained by Bonferroni conservative correction for total number of probes) was used to define significant associations of lipids with probe level gene expression.

Similarly, lipid residuals were used as dependent variable and average methylation value was used as the independent variable to test the association of lipids with methylation. The proportion of variance in lipids explained by the gene expression variation and methylation variance were estimated based on the log-likelihoods as implemented in the R package rcompanion ^50^. For most of the lipids, multiple gene expression probes were associated. Hence to avoid overfitting and multi-collinearity, we used penalized regression methods as implemented in glmnet of the R package caret ^51^ to reduce the number of probes in the regression model. The list of probes retained in the glmnet model was used to estimate the variance contributed by the gene expression.

For analysis of GWAM (Table 4), r^2^ is McFadden’s pseudo-r^2^. p-value for h^2^ is the p-value for test of significant additive genetic effects (h^2^=heritability). Thus p-value for h^2^ < 0.05 indicates significant heritability. Regression coefficients are based on average methylation at CpG sites excluding any with known SNPs influencing lipid levels.

### Lipid shorthand notation

Lipids are named according to the LIPID MAPS convention ^52^. Lipid abbreviations are as follows: ceramide (Cer), cholesterol ester (CE), diacylglycerol (DG), lysophosphatidylcholine (PC), phosphatidylcholine (PC), phosphatidylethanolamine (PE), phosphatidylinositol (PI), sphingomyelin (SM) and triglyceride (TG).

## Results

### Participant characteristics

Plasma lipidomics was performed on *n*=330 individuals, 260 of these were used for heritability analyses. Characteristics of the MZ (n=150, 100 females) and DZ (n=110, 79 females) twins with available plasma for heritability analyses are presented in Table 1. There were no group differences between MZ and DZ twins on these characteristics except in HDL-C levels, which were higher in MZ twins relative to DZ twins (p<0.05), but did not remain significant after correcting for multiple comparisons.

**Table 1.** Participant characteristics for heritability analyses.

|  | MZ (n=150) | DZ (n=110) | statistic | p-value |
| --- | --- | --- | --- | --- |
| Age | 75.7 (5.47) | 76.07 (5.31) | -0.548 | 0.584 |
| Females | 100 (67%) | 79 (72%) | 0.785 | 0.376 |
| Education (yrs) | 10.99 (3.18) | 11.2 (3.18) | -0.475 | 0.635 |
| BMI (kg/m<sup>2</sup>) | 27.934 (4.74) | 27.5 (4.92) | 0.776 | 0.438 |
| WHR | 0.89 (0.09) | 0.89 (0.08) | 0.164 | 0.87 |
| MMSE | 28.9 (1.37) | 28.95 (1.76) | -0.062 | 0.95 |
| LDL-C (mmol/L) | 2.77 (0.97) | 2.78 (0.97) | -0.078 | 0.938 |
| HDL-C (mmol/L) | 1.73 (0.46) | 1.60 (0.44) | 2.341 | 0.02 |
| Cholesterol (mmol/L) | 5.08 (1.01) | 4.98 (1.12) | 0.822 | 0.412 |
| Triglyceride (mmol/L) | 1.30 (0.54) | 1.32 (0.56) | -0.298 | 0.766 |
| *APOE* ε4 carrier* | 35 (26%) | 27 (28%) | 0.118 | 0.731 |

Means (SD) are presented for continuous variables, while n (%) is presented for categorical variables. Comparisons of MZ and DZ pairs used *t* tests for continuous variables and $\chi^2$ tests for categorical variables.
Abbreviations: MZ = monozygotic, DZ = dizygotic. body mass index (BMI), mini-mental state exam (MMSE), waist-hip ratio (WHR), low density lipoprotein cholesterol (LDL-C), high density lipoprotein cholesterol (HDL-C).
\*excludes participants with missing data (n=231 participants with *APOE* genotype data)

### Heritability

Heritability of lipids was computed using the classical ACE model. Classical lipid measures of total cholesterol, LDL, HDL and triglycerides were significantly heritable (*h*^2^=0.427, 95% C.I. = [0.075, 0.592], 0.404, 95% C.I. = [0.121, 0.573], 0.419, 95% C.I. = [0.027, 0.766], and 0.427, 95% C.I. = [0.181, 0.623] respectively). HDL had a substantial C component (i.e., common environment; *h*^2^C = 0.27, 95% C.I. = [0.00, 0.48]). For individual lipid species measured, 27 out of 203 (13.3%) were significantly heritable with a median heritability of *h*^2^ = 0.433, ranging from 0.287 for TG (18:0/17:0/18:0) to a maximum of 0.59 for Cer (d17:1/24:1).

The percentages of heritable lipids from the total pool of identified lipids in each lipid class is summarised in Figure 1A. Heritability estimates across lipid class and by individual lipid for significantly heritable lipids are summarised in Figure 1B and Table S1. Ceramides (Cer) had the highest heritability estimates (range *h*^2^=0.433 – 0.59), where 9 out of 20 species were significantly heritable. For triglycerides (TG), 12 of out 59 species measured were heritable (range *h*^2^= 0.287-0.495). Among diacylglycerols (DG), 3 species out of 10 were heritable (range *h*^2^=0.422-0.544). Only 3 phospholipids were heritable, including 2 of 58 phosphatidylcholines (PC) and 1 out of 5 phosphatidylethanolamines (PE), (range *h*^2^ = 0.327 – 0.413). Cholesteryl ester (CE), lysophosphatidylcholine (LPC), phosphatidylinositol (PI) and SM (sphingomyelin) species were not significantly heritable, with median heritability for non-significant lipids at *h*^2^=0.23, and near zero heritability for LPC species. Heritability estimates obtained for summed lipid groups (Table S2) were mostly similar to that of the individual lipids, though there were some differences. For example, the sum of monounsaturated SM species was heritable whereas no individual SM was significantly heritable. A complete heritability table for all lipids is presented in Data S1.

**Figure 1.**
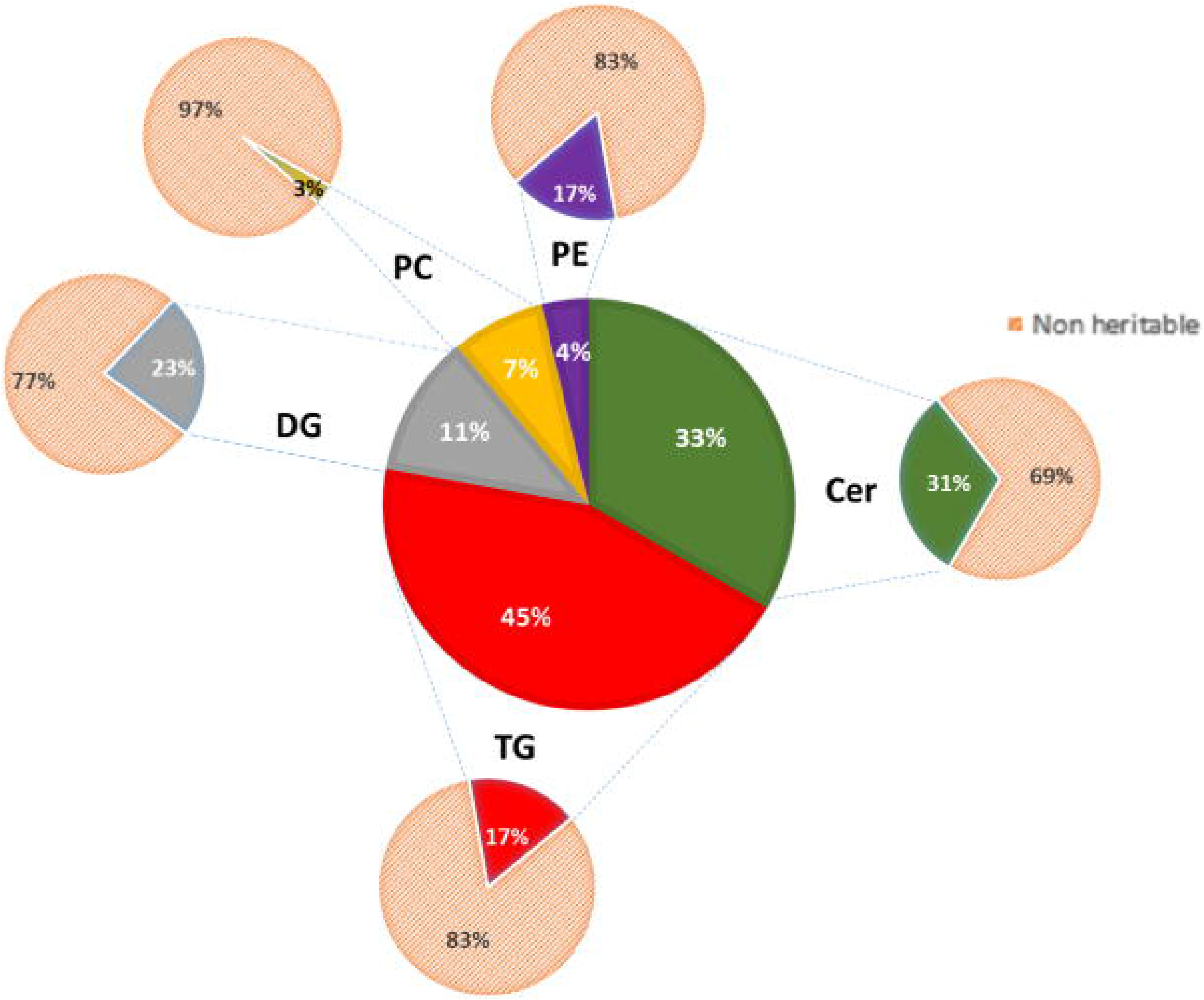
Heritability of lipids. (A) Percentage distribution of heritable lipids. The central wheel represents significantly heritable lipids and their percentage distribution by lipid class. Smaller wheels emanating from each sector represent proportions of these heritable lipids compared to total measured lipids of that class, such that the sum of these smaller wheels equals the total pool of 207 individual lipids measured. For example, 45% of significantly heritable lipids belonged to the TG lipid class, and these heritable lipids represented 17% of total measured plasma TG. (B) The distribution of heritability (h^2^), estimated from the ACE model, for each individual lipid species grouped according to class. A, C and E represent standardised variance components due to additive genetic (A=heritability), common/shared environment (C) and unique environment (E). Boxplots show median with interquartile range for each class. Dark circles represent heritable lipids, as opposed to grey circles, which represent lipids that were not significantly heritable. Minimum heritability is h^2^>0.287.

### Genetic, Environmental and Phenotypic Correlations

Genetic and environmental correlations were estimated for significantly heritable lipid species and lipid classes. Median genetic correlations between Cer species were high (r_g_=0.94), as were TG (r_g_=0.81) and DG (r_g_=0.73) species. DG and TG were also highly genetically correlated with each other (r_g_=0.70), as were Cer species with monounsaturated SM (r_g_=0.83). Median phenotypic correlations between Cer species, between TG species and between DG species were r_p_=0.85, 0.61, and 0.53 respectively, and r_p_=0.51 between TG and DG species, and r_p_=0.83 between Cer and monounsaturated SM. Median unique environmental correlations were moderately lower than corresponding genetic correlations (r_e_=0.75, 0.56 and 0.53 for Cer, TG and DG respectively, and r_e_=0.45 between TG and DG, and r_e_=0.72 between Cer and monounsaturated SM). Further, traditional lipids (LDL-C, HDL-C, total cholesterol and TG) had poor genetic and phenotypic correlations with individual lipid species, apart from traditional triglyceride measures, which was highly correlated with individual TG and DG species. A genetic correlation matrix heatmap is shown in Figure S1.

### Association with Gene Expression

The association of lipids (n=209) with probe level gene expression (n=35,971) was analysed using linear mixed models via the R package nlme ^49^. We found significant gene expression probe associations (n=3568) with 47 individual lipids (7 DG, 2 PC, 1 PE, 37 TG; see Data S2 and Data S6). Of these associations, 15 were linked to significantly heritable lipids (12 TG, 3 DG, n= 380 unique probes). In fact, we found that all significantly heritable TG and DG species were also significantly associated with gene expression of particular transcripts. An additional 32 individual lipids (25 TGs, 4 DGs, 2 PCs and 1 PE, n= 276 unique probes) without significant heritability were significantly associated with gene expression. In regards to traditional and grouped classes of lipids, there were also significant gene expression associations with HDL-C, total TG, and grouped TGs regardless of total carbon number or number of double bonds. No significant gene expression associations were identified for LDL-C. There was a modest but non-significant positive correlation between variance explained by gene expression of probes and heritability (p>0.05, Figure 2 and Data S2). This implies that gene expression accounts for some but not all the variance in heritable lipid levels.

**Figure 2.**
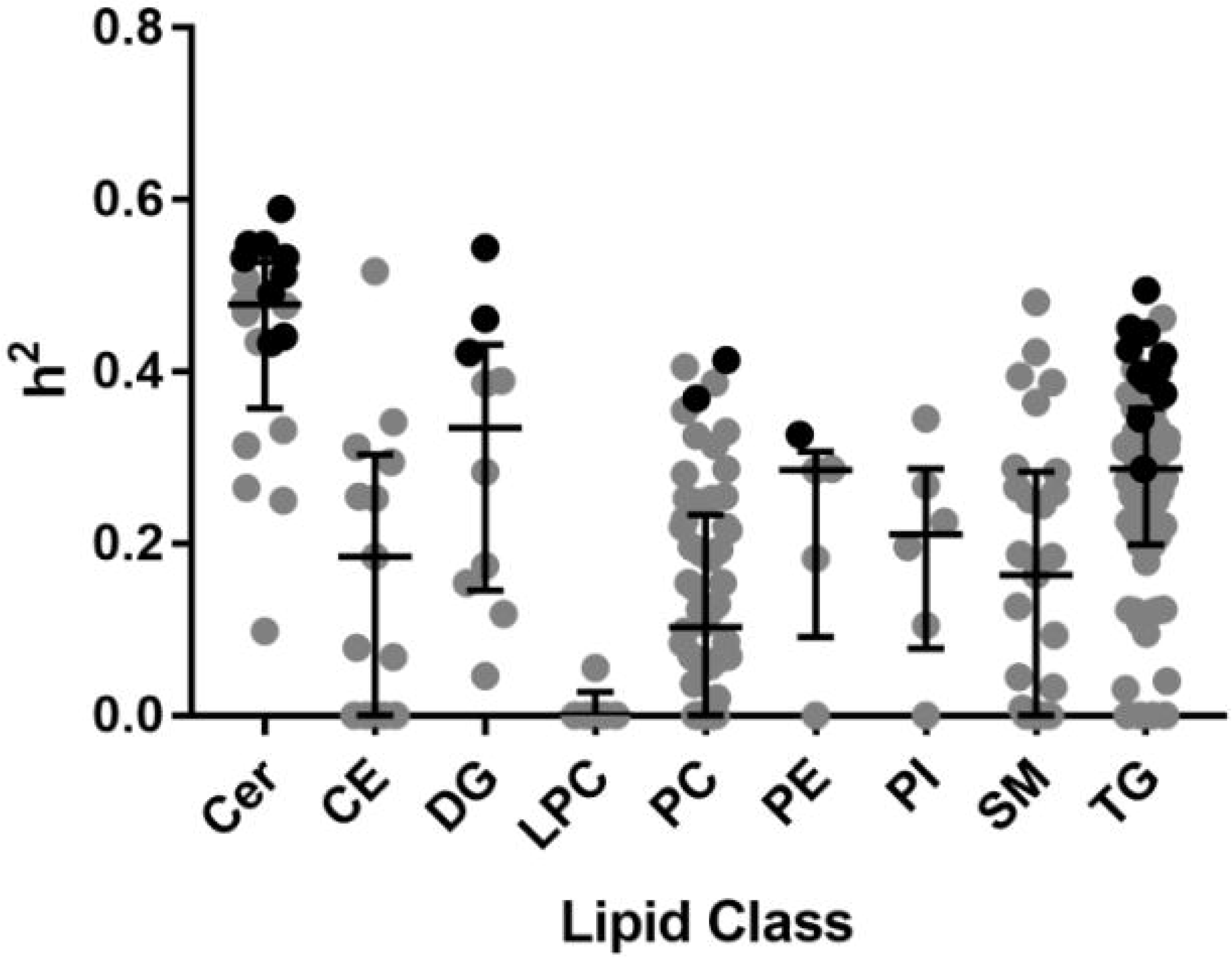
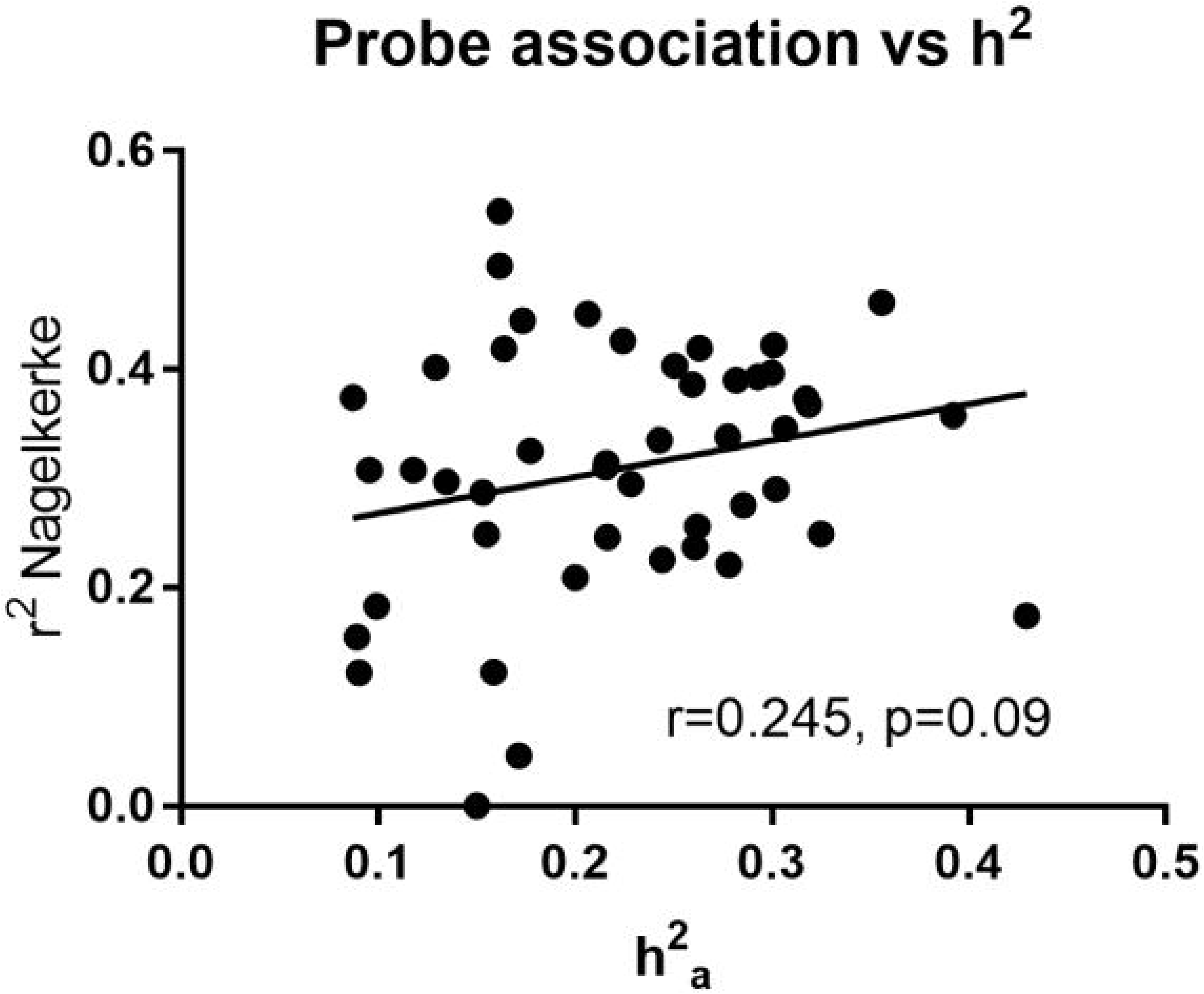
Heritability estimate (h^2^_a_) vs total variance explained (Nagelkerke r^2^) by gene expression probe transcripts for heritable lipids.

Since the bulk of significant gene expression associations were with TG, we examined the relationship of gene expression associations for TG species by degree of saturation, classifying each TG species as being saturated (no fatty acyl double bonds), monounsaturated (possessing one double bond), or polyunsaturated (possessing two or more double bonds). We then investigated how many transcripts were associated with a low, medium and high number of lipids, by counting the number of gene transcripts significantly associated with either 1-2 lipids, 3-8 lipids, and over 8 lipids in that class (in the case of polyunsaturated TG). Generally, only a few gene transcripts were associated with many lipids, regardless of saturation level. There were 282 gene transcripts associated with 1-2 TGs in the saturated TG class, but only 6 were associated with at least three different TGs in that class.

Table 2 summarises the number of significantly associated gene transcripts among each TG saturation class, while Figure 3 is a Venn diagram identifying gene transcripts that are unique or shared across saturation classes for significantly heritable TG lipids (Figure 3A) and non-heritable TGs only (Figure 3B). The total list of gene transcripts associated with lipids can be found in Data S3 and S4, while Data S5 and Data S6 show gene transcripts ordered by TG degree of saturation and total number of carbons. For example, ribosomal protein L4 pseudogene 2 (*RPL4P2*), A disintegrin and metalloproteinase domain-containing protein 8 (*ADAM8*) and Adipocyte Plasma Membrane Associated Protein (*APMAP*) were uniquely associated with saturated TG when considering heritable TG lipids.

**Table 2.** Gene expression associations among TG lipids.

| TG Class | Number of Associated Lipids | Number of Transcript Associations |
| --- | --- | --- |
| Saturated TG | 1-2 | 282 |
|  | 3-8 | 6 |
| Monounsaturated TG | 1-2 | 59 |
|  | 3-8 | 7 |
| Polyunsaturated TG | 1-2 | 243 |
|  | 3-8 | 119 |
|  | >8 | 9 |

Note. Table 2 lists number of gene expression associations common to a maximum of 1-2, 3-8 and >8 lipids in each TG saturation class (saturated, monounsaturated, and polyunsaturated TG).

**Figure 3.**
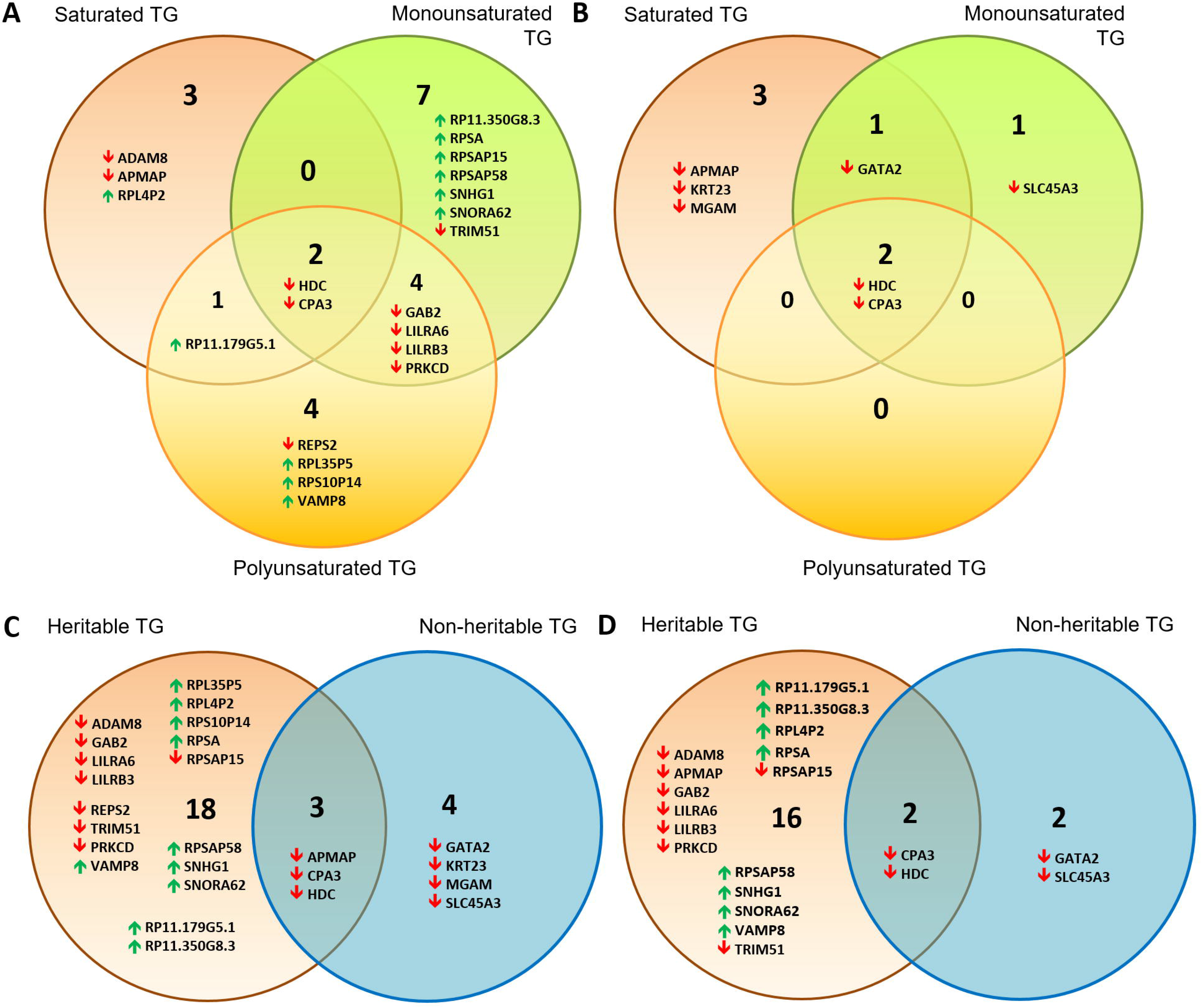
Venn diagrams showing distribution of gene transcripts associated with a majority of TG lipids. These were subdivided into those associated with saturated vs monounsaturated vs polyunsaturated lipids for (A) significantly heritable TGs and (B) non-heritable TGs. Also shown are heritable vs non-heritable set of significant gene expression associations of TG lipids for (C) double bond group/saturation (Data S5) and (D) total number of carbons (<49 carbons, 49-55 carbons and 56+ carbons, Data S6). Gene transcripts included in these Venn diagrams were those significantly associated with the highest and second highest number of lipids of a particular saturation class (A and B), or among heritable and non-heritable lipids (C and D). Upwards and downwards arrows indicate positive and inverse gene expression associations with lipid levels respectively.

Interestingly, there were a number of transcripts associated with a maximum of 1-2 TG lipids (e.g. 1-2 saturated lipids had 282 hits). In a majority of cases, these associations were driven by a specific TG lipid (among saturated TGs, this was TG(16_0/16_0/24_0), among monounsaturated TGs, this was TG(16_0/14_0/18_1) and for polyunsaturated TGs, these were TG(19_1/18_1/18_2), TG(16_0/18_1/23_1), TG(16_0/22_6/22_6) and TG(25_0/18_1/18_1)). These lipids tended to have a medium to high total carbon count (i.e. >55 carbons). By contrast, our analysis also found gene expression of histidine decarboxylase (*HDC*) and carboxypeptidase A3 (*CPA3*) to be significantly associated with all TGs irrespective of the number of total carbons and number of double bonds. In fact, *HDC* and *CPA3* were also significantly associated with other lipids including DG and HDL-C (Data S6). Notably, there were some differences between the gene transcript association profiles of significantly heritable vs non-heritable lipids; many more gene transcript associations were unique to heritable as opposed to non-heritable TGs (Figure 3A-D, Table S3). For example, pseudogenes appearing in the heritable lipid list do not appear in the non-heritable list. Comparing transcribed genes associated with TG lipids by total number of carbons (<49 carbons “low”, 49-55 carbons “medium” and 56+ carbons “high”) also yielded a similar outcome (Figure 3D).

Furthermore, the majority of transcriptome associations with non-heritable lipids were inverse associations, whereas the lipid-transcriptome associations for heritable lipids were a mix of positive and inverse associations, suggesting a diverse impact of these lipids on cellular function. It is also interesting that the majority of inverse lipid-transcriptome associations encode protein coding transcripts (15/17 total), and only 2/17 were non-protein coding RNAs/pseudogenes. By contrast, the majority of positive lipid-transcriptome associations were for non-protein coding pseudogenes (9/11) and only 2/11 were protein coding.

### Functional pathways of associated gene transcripts

The majority of the protein coding transcriptome which associates with our lipidomic data has some association with inflammatory and vascular pathways (Table 3), with possible roles in the central nervous system (CNS). The STRING and BioGRID databases ^53; 54^ were used to provide functional information on genes identified in the lipid-transcriptome analysis. Some other notable pathways include vasoactive peptides, vesicular transport and pseudogenes/non-protein coding genes.

**Table 3.** Functions of genes with significant lipid-gene transcriptome associations.

| Biological Pathways | Gene Transcripts* | Relevance to the CNS |
| --- | --- | --- |
| **Inflammation** |  |  |
| Innate immunity | ↓*LILRB3*, ↓*MGAM* |  |
| Adaptive immune response | ↓*LILRA6* |  |
| Host Defense | ↓*FPR1*, ↓*TRIM51* | FPR1 found in neural glial cells, astrocytes and neuroblastoma <sup>55</sup>. |
| Allergic Response | ↓*ADAM8*, ↓*HDC*, ↓*CPA3* | ADAM8 may regulate cell adhesion during neurodegeneration <sup>56</sup>.<br>HDC as a histidine decarboxylase, produces histamine, which in the CNS is a neurotransmitter <sup>57</sup>. |
| Class I MHC antigen binding | ↓*LILRA6*, ↓*LILRB3* |  |
| B-Cell response/receptor signalling | ↓*GAB2*, ↓*LILRB3*, ↓*PRKCD* | GAB2 is associated with Alzheimer's disease. By activating PI3K, increases amyloid production and microglia-mediated inflammation. Several *GAB2* SNPs are associated with late-onset Alzheimer's disease <sup>58</sup>. |
| Mast Cell Degranulation | ↓*CPA3*, ↓*HDC*, |  |
| **Vasoactive Actions** |  |  |
| Regulation of vasoactive peptides (e.g., endothelin, angiotensin 1, snake toxins, etc) | ↓*GATA2*, ↓*CPA3*, |  |
| Epithelial Cell Integrity | ↓*KRT23*, ↓*PRKCD* |  |
| Cell Adhesion | ↓*APMAP* | APMAP supresses brain Aβ production <sup>59</sup>. |
| DNA Regulation | ↑ *RPSA*, ↑ *SNORA62*, ↑ *SNHG1* |  |
| Vesicle/Endosome Regulation/Transport | ↑ *VAMP8*, ↓*REPS2*, ↓*SLC45A3* | SLC45A3 regulates oligodendrocyte differentiation <sup>60</sup>. |
| Pseudogenes/non-protein coding | ↓*S100A11P1*, ↓*RPSAP15*,<br>↑ *RP11-179G5.1*,<br>↑ *RP11-350G8.3*,<br>↑ *RPL35P5*, ↑ *RPL4P2*,<br>↑ *RPS10P14*, ↑ *RPSAP15*,<br>↑ *RPSAP58*, ↑ *SNHG1*,<br>↑ *SNORA62* | Regulatory roles. Gene silencing, affects mRNA stability. |

Directions of arrows indicate either positive (upwards facing) or inverse (downwards facing) lipid-gene transcriptome associations. Even though our transcriptomic data was for the blood transcriptome, some of these genes also have functions in the CNS or associations with neurodegenerative diseases (far right column).

### Association of DNA methylation levels at specific CpG sites with lipid and gene expression

To gain insight into the relationships between lipid levels and DNA methylation of CpGs at specific genes, we selected gene transcripts significantly associated with lipids and identified associations between DNA methylation at CpG sites within close proximity to these gene transcripts, and lipid expression (Data S7). We found significant associations of DNA methylation (p<0.05) with four lipids: PE(16:0_20:4), TG(25:0_16:0_18:1), TG(18:0_17:0_18:0) and TG(18:1_18:2_18:2). Of these, two were heritable - TG(25:0_16:0_18:1) and TG(18:0_17:0_18:0).

We also examined the relationship between gene expression and DNA methylation at specific CpG sites of genes whose transcripts were associated with significant heritability (Data S8). We found 19 significant CpG site-gene expression associations related to four unique lipids (TG(19:1_18:1_18:2), TG(15:0_16:0_18:1), PC(20:2_18:2), TG(16:0_18:1_23:1), but these associations were a very minor subset of all CpG site-gene expression associations. Therefore, we did not find sufficient evidence to suggest that DNA methylation at specific CpG sites drives changes in gene expression, though we acknowledge this analysis lacks sufficient power to be conclusive.

### Association of lipids with genome wide average DNA methylation (GWAM)

We then explored associations of genome wide average methylation with lipid levels and found significant associations of all five LPCs (and the total LPC sum) with GWAM (range beta = −0.22 to - 0.27, see Table 4). Notably, four TGs were also significantly inversely associated with GWAM (beta = −0.18 to −0.23). Further, only two other lipids were positively associated with GWAM, namely one CE and one PC (beta = 0.21, and 0.18 respectively, Table 3). None of these lipids was significantly heritable, with maximum heritability of 0.39, though one TG (TG18:1_17:1_22:6) was borderline significant (p=0.05 for h^2^), with a maximum of two significant gene expression associations (for TG18:1_17:1_22:6 and TG18:1_20:4_22:6).

**Table 4.** Regression of lipid residuals significantly associated with genome wide average DNA methylation levels

| Lipid | Beta | SE | t | p-value | $h^2$ | p-value for $h^2$ |
| --- | --- | --- | --- | --- | --- | --- |
| CE(20:3) | 0.21 | 0.09 | 2.34 | 2.31E-02 | 0.31 | 0.30 |
| LPC(15:0) | -0.22 | 0.09 | -2.54 | 1.39E-02 | 6.51E-16 | 1 |
| LPC(16:0) | -0.27 | 0.09 | -3.12 | 2.90E-03 | 3.82E-14 | 1 |
| LPC(17:0) | -0.21 | 0.09 | -2.34 | 2.30E-02 | 2.82E-14 | 1 |
| LPC(18:1e) | -0.21 | 0.09 | -2.44 | 1.81E-02 | 3.52E-17 | 1 |
| LPC(26:0) | -0.27 | 0.09 | -3.10 | 3.07E-03 | 0.056 | 0.87 |
| PC(39:3) | 0.18 | 0.09 | 2.12 | 3.84E-02 | 0.39 | 0.14 |
| TG(18:1_17:1_22:6) | -0.18 | 0.09 | -2.05 | 4.51E-02 | 0.31 | 0.05 |
| TG(18:1_18:1_22:5) | -0.23 | 0.09 | -2.69 | 9.58E-03 | 3.42E-15 | 1 |
| TG(18:1_20:4_22:6) | -0.21 | 0.09 | -2.41 | 1.96E-02 | 2.98E-15 | 1 |
| TG(19:0_18:1_18:1) | -0.18 | 0.09 | -2.14 | 3.73E-02 | 0.312 | 0.29 |
| GroupLPC | -0.24 | 0.09 | -2.74 | 8.32E-03 | 1.88E-15 | 1 |

Notes. Associations of GWAM with lipid residuals (adjusted for age, sex, education, BMI, lipid lowering medication, smoking status, experimental batch and *APOE* $\epsilon 4$ carrier status).

## Discussion

### Heritability Estimates

In this study, we evaluated the relative contributions of genetic versus environmental factors to the plasma lipidome among older Australian adults aged 69-93 years. As hypothesised, both genetic and environmental factors contribute to shaping the plasma lipidome, though in our sample of older individuals, environmental factors were predominant, with only 13.3% of individual lipids analysed being significantly heritable. The median heritability of heritable lipids was h^2^ = 0.433, indicating a moderate level of heritability which compares well with an estimate of 36.2% provided for metabolites from a genome-wide genotyping study in subjects aged 60 years and over ^61^. The effect of common environment (C) was minimal for all lipid measures, except for a low to moderate finding for HDL-C (0.27), which is consistent with previous studies ^12; 62^, indicating that the shared environment early in life is an important contributor to HDL-C variance later in life.

Traditional lipid measures of LDL-C, HDL-C, total cholesterol and TG were significantly heritable, consistent with previous studies ^12; 13^, though our estimates for these traits (range 0.40-0.47) were lower than estimates from other studies, where heritabilities have been reported to exceed 0.60 ^12; 63^. Interestingly, one of these studies compared heritability using data from three cohorts around the world, and found heritability estimates of these traits among Australian twins to be lower than the same estimates in Dutch and Swedish twin pairs ^63^. It is likely that heritability differences between different studies are a product of differences in ethnicity, cohort and age, leading to substantial variance in reported heritabilities from study to study.

Our analyses also indicated that there is differential heritability depending on the lipid class examined. While no individual class had over 50% of its lipids significantly heritable, the largest proportion of heritable lipids was found for TG, followed by Cer and DG. By contrast, only three phospholipids were significantly heritable, two of these being PCs (out of a total of 58 PCs assessed), and one PE. None of the remaining classes examined (SM, CE, LPC and PI) yielded significantly heritable lipids, with LPCs having virtually no heritability in this study. Further, it is important to note that the pattern of heritability across summed lipid traits often did not match the heritability of individual lipids in a given lipid class or subclass, likely owing to the broad range of heritability estimates obtained, which was also reported in a previous study ^31^. Thus, for heritability analyses, it appears important to analyse heritability estimates for individual species as opposed to summed lipid classes.

The high heritability of Cer in this study supports data from family-based GWAS which found 36 ceramides to be significantly heritable, with heritability estimates as high as 0.63 ^64^. One recently published German twin study using data from *NutriGenomic Analysis in Twins* (NUGAT), examined the extent to which lipidomic changes in response to a high fat diet intervention are heritable and yielded a similar range of heritabilities for individual lipid species, with estimates ranging from 0-62% ^31^. This study identified 19 of 150 plasma lipid species to be highly heritable (h^2^>0.40), which is not dissimilar to the number of significantly heritable individual lipid species identified in the present study (27 of 207). However, the heritability of various classes often did not corroborate our findings. For example, in the NUGAT study, LPC and PE were reported to be moderately heritable (0.25<h^2^<0.35), while SMs had high heritability, as opposed to ceramides which were reported to be lowly heritable. By contrast, our study found high heritability of ceramides with no significantly heritable SMs and virtually zero heritability of LPCs. One possible explanation for these differences is that heritabilities may change across the lifespan ^13^. Age-dependent heritability has been reported for LDL-C and HDL-C ^13^, and also in BMI, where heritability estimates are lower in older adults compared to young adults ^65^. The age range of NUGAT participants was 18 to 70 years, with a median of 25 years, whereas the OATS cohort consisted of much older individuals ranging from 69 to 93 years. This could be especially important considering the potential impact of pre- and post-menopausal status on lipid profiles ^66; 67^ among women, who comprise a majority of participants in both OATS (n=179, 68.8%) and the NUGAT study (n=58, 63%). Additionally, the NUGAT study features a substantially smaller sample size of 46 twin pairs (34 MZ and 12 DZ twin pairs, vs. 75 MZ and 55 DZ twin pairs in the present study) and NUGAT heritabilities were based on linear mixed models with an additional unknown effects variance added. Thus it may be more difficult to ascertain the C component of the NUGAT study, which we found to be negligible for all lipids except for HDL. A more recent publication of a Finnish population based study (FINRISK) reported SNP-based heritability of lipid species to be in the range 0.10-0.54 ^68^, and found Cer to be the most heritable species, corroborating findings from the present study, though heritability of some other lipid classes, such as LPC was markedly higher than reported in the present study. As with the NUGAT study, some differences could be attributable to the younger age range of participants (25 – 74 years) in FINRISK relative to OATS, and potentially the low sample size for SNP-based heritability calculations.

### Genetic correlations

High within-class genetic correlations between individual Cer, TG, and DG species (all r> 0.70) suggest similar genetic influences between lipids of the same class. Further, Cer species and monounsaturated SM also exhibited high genetic correlations, as did TG and DG. Metabolically, Cer and SM belong to the sphingolipid class where SM can be converted to Cer via sphingomyelin phosphodiesterase ^69^, while TG and DG are interconvertible, where TG can be metabolised to DG by adipose triglyceride lipase (ATGL), or DG to TG through the addition of acyl CoA via DG acyltransferase (DGAT) ^70^. Further, genetic correlations were all higher than the corresponding environmental correlations, indicating heritable lipids of similar class have a strong shared genetic basis relative to the unique environment. Our results suggest that the heritable lipidome is regulated by overlapping genes which are associated with multiple lipids, especially lipids that belong to the same class, or are related by a connected metabolic pathway. Nevertheless, environmental correlations were still high for these lipids suggesting the importance of environmental factors on lipid levels. Traditional lipids (total triglyceride, LDL-C, HDL-C and total cholesterol) had low genetic and phenotypic correlations with individual lipid species, except for triglyceride measures, which were highly correlated with TG and DG species. This finding confirms previous results ^68^ and suggests some differences between variance in traditional lipid measures and variance in the lipidome at the individual lipid species level.

### Lipid-Transcriptome Associations

Transcriptome associations of both heritable and non-heritable triglycerides, which represented the largest component of our lipidomics dataset, were assessed. We anticipated that both heritable and non-heritable lipids would have gene transcript probe associations, since endogenous triglycerides are derived from essential dietary fatty acids, such as linoleic acid, or other fatty acids substantially derived from dietary sources (such as linolenic acid and docosahexaenoic acid). Gut microbiota (microbiome) can also have an effect on the dietary lipidome, prior to absorption, representing another “environmental” contributor, to lipid abundance and structure ^71^. Once absorbed, the environmentally sourced lipid milieu becomes available for genetically regulated structural change by a diversity of lipid modifying machinery. This includes families of elongase and desaturase enzymes responsible for modifying fatty acid chain length and saturation level ^72^, as well as a plethora of synthetases which assemble complex lipids such as the triglycerides and phospholipids ^73^. Interestingly, in our gene/lipid transcriptomic association list (Tables 3 and 4/Figure 3 and Figure 4), such structure regulating genes do not appear. Instead, the transcripts reveal genes which regulate other physiological and cellular functions, particularly those involved with immune and vascular functions (Table 4). We also found an upregulation of pseudogenes, which could play important regulatory roles, such as in gene silencing ^74^.

**Figure 4.**
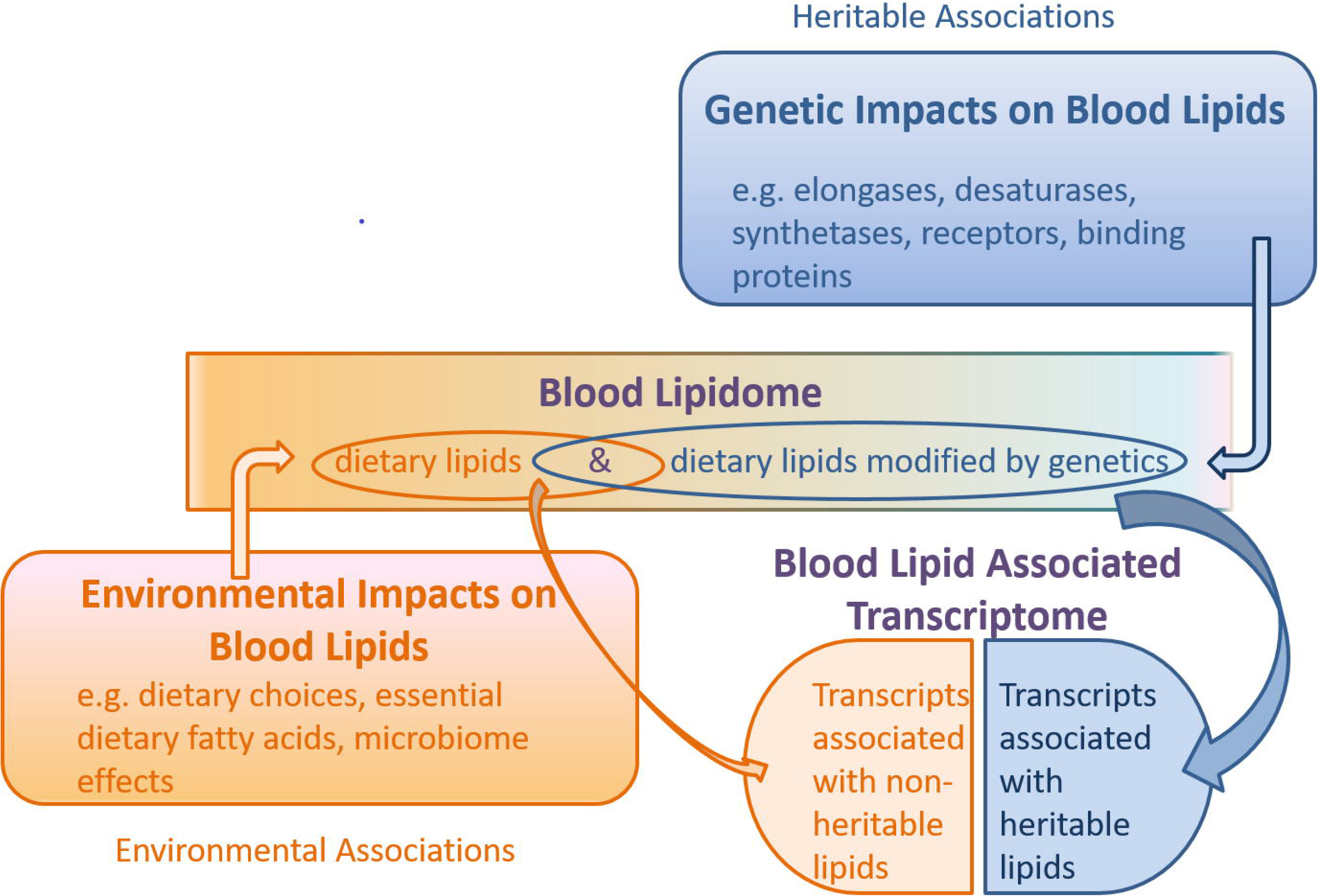
Schematic of the combined genetic and environmental influences on the blood lipidome, and the association of this lipidome with the blood transcriptome. Under this model, non-heritable lipids could affect gene transcription, while heritable lipids could also affect gene transcription (collectively “blood lipid associated transcriptome”), but are possibly modified upstream by genetic machinery such as elongases, desaturases, synthetases, receptors and binding proteins. Gene transcripts encoding these enzymes and proteins may be independent of the “blood lipid associated transcriptome” noted in this study.

From this, we infer that the genes which are thought to account for the substantially heritable phenotype of our triglyceride group (i.e. via lipid metabolic processes) are not necessarily the same as those reflected in the lipid-transcriptome associations. This might be the case if the heritable aspect of our lipid list is driven by lipid modifying genes (such as desaturases, elongases, fatty acid synthases and synthetases), while the blood transcriptome is associated with the endogenous lipidome, which is a product of both environment and genetics (a feedback loop of sorts). We model this hypothesis in Figure 4. This also complements our finding that variance in lipid levels due to heritability is only partially accounted for by gene expression of associated transcripts.

### Biological effects of the lipid associated blood transcriptome

Our lipid-transcriptome analysis revealed strong associations of lipids with gene transcripts involved in modulating immune and vascular function. Interestingly, a previous twin study found a minor subset of the immune system is modulated by genetic influences, such as the homeostatic cytokine response ^75^, and many of the associated gene transcripts in the current study including Solute carrier family 45 member 3 (*SLC45A3*)*, CPA3* and *HDC* were previously reported in a study of lipid and immune response ^76^. Thus, some of the transcriptome associations uncovered could reflect lipid-modulated innate immune responses. Since this protein coding transcriptome has largely negative associations with lipid levels, we infer that it is moderating/suppressing inflammation or adverse vascular events. On the other hand, high fat diet in mouse models leads to elevated gene transcription related to white adipose tissue and liver metabolism, and after a prolonged high fat dietary regimen, activation of inflammatory pathways ^77^. We postulate that lipid levels are normally linked to the suppression of inflammatory responses to maintain homeostasis, but become associated with activation of inflammatory responses following metabolic overload, such as in diabetes mellitus or obesity ^78; 79^. Indeed, the authors of this study only noted significant upregulation of genes associated with inflammatory pathways after six weeks of high fat diet consumption, in contrast to genes associated with lipid metabolism, which were upregulated directly following a high fat diet.

We found most of the associated lipid-protein coding transcriptome to be membrane proteins, which suggests a possible interaction between lipids and protein function at the cellular surface. This would also explain transcripts being associated with proteins involved in phosphorylation and other signalling pathways. Vesicle associated membrane protein 8 (VAMP8) is involved in cellular fusion and autophagy. A couple of transcripts are associated with endothelial function. CPA3 is involved in anti-proteotoxic effects by proteolytically cleaving peptides with potentially harmful physiological effects such as vasoconstriction peptides (endothelin and angiotensin 1) and snake venom peptides. Other identified transcripts are involved with production of vasoactive peptides. GATA2 regulates endothelin-1 gene expression in endothelial cells, and PRKCD phosphorylates ELAV Like RNA Binding Protein 1 (ELAVL1) in response to angiotensin-2 treatment ^80^.

### Lipid associations with DNA Methylation

To assess possible mechanisms contributing to variance of non-heritable lipids, we compared average DNA methylation levels over 450,000 different DNA methylation sites among MZ twins. DNA methylation is a well characterised epigenetic mechanism by which a gene expression profile can be regulated and inherited independent of the genetic sequence ^81^, and involves the addition of a methyl group (-CH_3_) to the base cytosine of 5’-cytosine-phosphate-guanine-3’ (CpG) dinucleotides ^82; 83^. Methylation of CpG clusters around promoter regions of genes typically leads to suppression of gene transcription. DNA methylation analysis revealed a suggestive level of significance for the association of GWAM with 4 TGs, 1 PC, and all 5 LPCs. None of these lipid species was significantly heritable, with the exception of TG(18:1_17:1_22:6), which was borderline heritable (p=0.05). In particular, all five LPCs and their summed total, which were extremely non-heritable (near zero), were also significantly associated with GWAM. Although only a small subset of lipids showed significant associations with GWAM (just 8 individual lipids of 180 non-heritable lipids), these findings do suggest that epigenetic factors such as DNA methylation could explain some of the variation associated with non-heritable lipids, especially very lowly heritable phospholipids and LPC, the least heritable lipid class in our data-set.

In previously published work, DNA methylation has been associated with environmental changes in lipid levels. Maternal lipids, passing from mother to child *in utero* at 26 weeks of gestation, lead to DNA methylation changes in the newborn ^84^. The lipids associated with DNA methylation changes included phosphatidylcholine and lysolipids – phospholipid degradation products. The authors hypothesised that the choline source from these lipid products could be important precursors for DNA methylation. Further, the direction of change was largely negative, with higher lipid metabolites associated with lower methylation levels of genes involved in prenatal development. While the association of LPCs with DNA methylation has not previously been identified, it is worth noting that LPCs are a major source of polyunsaturated fatty acid (PUFA) for the brain ^85^ and regulate gene transcription through sterol regulatory-element binding protein (SREBP) pathways ^86^. Thus, LPC is an important lipid to convey dietary sources of PUFAs into the brain and regulate gene transcription.

These findings add to studies conducted in animal models which also show that nutrients taken by the mother are passed on to offspring during pregnancy, and may have a lasting impact on gene expression through DNA methylation ^87^. Dietary restriction has also been shown to attenuate age-related hypomethylation of DNA in the liver, resulting in the downregulation of genes involved in lipogenesis and elongation of fatty acid chains in TGs, leading to a shift in the TG pool from long chain to medium and shorter chain TGs ^88^. In summary, there is evidence to suggest that lipids can influence DNA methylation levels, while genes related to lipid metabolism can also be regulated in response to DNA methylation.

Interestingly, when we attempted to focus on DNA methylation at specific CpG sites within close proximity to genes whose transcripts were significantly associated with lipids, we found a few associations with lipids and with gene expression, but little overall evidence to indicate that DNA methylation drives gene expression of these transcripts. More work needs to be done to clarify these relationships using a larger sample size.

### Limitations and future perspectives

There are some important limitations to this work. Firstly, this study covers a fairly wide age range in older aged adults (69-93 years). Very few heritability studies have focused on the lipidome in this age bracket. It is thereby important to stress the findings of this study may not necessarily generalise to the whole population and could be unique to the elderly, specifically those aged over 70 years. We suspect that in this cohort, environmental factors would dominate given the time in which these exposures are allowed to accumulate and shape the lipidome. Some of the heritabilities reported may vary longitudinally, owing to the dynamic contribution of genetic and environmental factors, and their interaction, across the lifespan [81]. In particular, heritability estimates may decrease where unique environmental exposures accumulate with time and become a dominant force in lipid modulation. By contrast, heritabilities may also increase where certain genes become more active in older age to shape a given phenotype. Given the age range of the cohort used, the results from the present study likely reflect a combination of both genetic and environmental influences on variation in the lipidome relevant to older age, and may provide important clues as to lipids and genes important in longevity. Some of these influences may underlie metabolic and lipidomic signatures previously described in very old individuals ^19; 42; 89; 90^. It is also important to emphasise that heritability estimates only represent the relative contribution of genetic and environmental influences. A “low heritability” score does not necessarily imply that there are no additive genetic effects, but rather that variation in the lipid profile among twins is largely mediated by the shared or unique environment. Further, we acknowledge that though we have included as many participants as possible from this study, there may be insufficient power to make substantive conclusions. Nevertheless, we believe our findings to be a good starting point for further investigation.

Transcriptomics data obtained through the Illumina microarray provides a broad overview of many potential gene transcript associations with measured lipids from the same individuals. However, these data were obtained using RNA from blood cells, which presents potential biases in the types of associations uncovered and could account for some of the immune regulatory genes uncovered. Nevertheless, given the strict cutoff p-value employed in the analyses, it is likely these associations reflect true roles of these lipids in immune function, and the genes we uncovered have previously been identified in other lipid-transcriptomic studies ^76^. We must emphasise that the transcriptome is influenced by many independent factors up- and downstream. The relationship between genetic variance (heritability) and the transcriptome is not clearcut. Nevertheless, we find some evidence that the transcriptome is linked to heritable plasma lipids and may explain a small proportion of their heritability. Additionally, while ceramides were the most heritable lipids, there were no significant gene expression associations with these lipids. This could be due to very low endogenous expression of ceramide synthases in leukocytes ^91^, though this pattern may be different in tissues where the most abundant CerS, CerS2, is highly expressed ^92^, such as in the kidney or liver ^91^.

Another major limitation is the fact that only average levels of DNA methylation (i.e. GWAM) were considered when associating with lipids, rather than DNA methylation at specific sites. This approach was necessary in order to avoid multiple testing correction for over 450,000 CpG methylation sites. The result is that the associated lipids showed at best suggestive significant associations with DNA methylation. The associations that we did find were for non-heritable lipids only, especially the least heritable LPCs, and were largely inverse. It is likely that based on previous studies, more significant associations with DNA methylation sites could be determined using greater selectivity of methylation sites at certain genomic regions. Further, as our analysis only showed that a small subset of non-heritable lipids were associated with GWAM, there is a still a lot variation in the lipidome not accounted for. CpG site specific analysis for particular genes did not find a relationship between DNA methylation and gene expression of these transcripts, though this analysis may lack power to detect these relationships. Other epigenetic mechanisms such as histone modification and chromatin structural changes could be implicated in regulating lipid metabolism, but are beyond the scope of this study. In spite of these limitations, this study provides a strong first step towards understanding some of the complex contributions of genes and the environment to shaping the human plasma lipidome at a lipid species level, especially among older individuals.

## Conclusion

In our study of older Australian twins combining lipidomics, transcriptomics and DNA methylation data, a small subset of plasma lipids was heritable and included largely Cer, TG and DG species. Most phospholipids, especially LPCs, were not significantly heritable. Significantly heritable lipids exhibited high genetic correlations between individual Cer, TG and DG species, as well as between Cer and SM, and between DG and TG, indicating shared genetic influences between lipids of the same class or metabolic pathways. Heritable lipids, especially TGs and DGs, were associated with a greater degree of gene transcript probe associations relative to the non-heritable lipids, and these transcripts were related to immune function and cell signalling rather than lipid metabolism directly. Thus, genes not related to lipid metabolism may still be associated with plasma lipid levels. Finally, associations of genome-wide average DNA methylation with highly non-heritable lipids, especially LPCs, suggest a potential mechanism by which environmental influences on lipids are conveyed. Overall, this study shows that a vast majority of plasma lipids are controlled by the environment, and hence modifiable, with genetic control still a major contributor to Cer, DG and TG lipid levels. Further, our study suggests a complex interaction between lipids, environment, DNA methylation and gene transcription.

## Acknowledgements

We acknowledge the contribution of the OATS research team (https://cheba.unsw.edu.au/project/older-australian-twins-study) to this study. The OATS study has been funded by a National Health & Medical Research Council (NHMRC) and Australian Research Council (ARC) Strategic Award Grant of the Ageing Well, Ageing Productively Program (ID No. 401162) and NHMRC Project Grants (ID 1045325 and 1085606). We thank the Rebecca Cooper Medical Research Foundation for their research support. We thank the participants for their time and generosity in contributing to this research. We would also like to acknowledge Ms. Mahboobeh Housseini’s assistance with OATS plasma biobanking.

This research was facilitated through Twins Research Australia, a national resource in part supported by a NHMRC Centre for Research Excellence Grant (ID: 1079102).

## Web Resources

The STRING v11.0 database and Biological General Repository for Interaction Datasets (BioGRID) were used to identify known and potential functional connections between gene-coded proteins that are associated with heritable lipids. STRING (https://string-db.org/) and BioGRID (https://thebiogrid.org) are free open-access resources available online.

