## Supplemental Information for "Genetic and environmental determinants of variation in the plasma lipidome of older Australian twins"

### Supplemental Methods

- **LC MS/MS quality control samples**

A set of quality control "standards" were injected every 20 runs. These controls included: (i) blank, to check for column and chromatography background levels, (ii) internal standards only, to check on system performance across a long sequence of runs, (iii) quality control plasma, to check for between run performance and enable calculation of between run and within run assay CV%.

- **Lipidsearch v4.1 search parameters**

We performed search on raw files using the databases "General" and "labelled standards". For peak detection, recalc isotope was set to "ON", RT interval = 0.0 min. We used product search for LC-MS method and the precursor and product tolerances were set at 5.0 ppm and 8.0 ppm respectively. The intensity threshold was 1% parent ion, and the m-score threshold was set to 2.0. For quantitation, mz tolerance was set at -5.0 ppm to 5.0 ppm, and the retention time range was set at -0.5 to 0.5 min. The m-score threshold was 5.0, and all lipid classes were selected for inclusion. Ion adducts included +H, +NH4 for positive ion mode.

#### Table S1. Heritable Lipid Species

| *Trait* | *$h^2$ (95% C.I.)* | *$h^2_c$ (95% C.I.)* | *$h^2_E$ (95% C.I.)* | *ICC MZ (95% C.I.)* | *ICC DZ (95% C.I.)* | *p-CE* | *p-AE* | *p-E* |
|---|---|---|---|---|---|---|---|---|
| Cer(d16:1_24:1) | 0.53 [0.22, 0.68] | 3.03E-14 [0.00, 0.23] | 0.47 [0.32, 0.66] | 0.53 [0.34, 0.68] | 0.27 [0.17, 0.38] | 0.005 | 1.00 | 8.30E-06 |
| Cer(d17:1_24:1) | 0.59 [0.28, 0.72] | 1.42E-15 [0.00, 0.24] | 0.41 [0.28, 0.58] | 0.59 [0.42, 0.72] | 0.29 [0.21, 0.42] | 0.002 | 1.00 | 1.97E-07 |
| Cer(d18:0_22:0) | 0.51 [0.18, 0.66] | 4.38E-16 [0.00, 0.26] | 0.49 [0.34, 0.67] | 0.51 [0.33, 0.66] | 0.26 [0.16, 0.39] | 0.008 | 1.00 | 7.05E-06 |
| Cer(d18:0_24:0) | 0.43 [0.01, 0.59] | 9.96E-17 [0.00, 0.32] | 0.57 [0.41, 0.76] | 0.43 [0.24, 0.59] | 0.22 [0.12, 0.37] | 0.045 | 1.00 | 2.62E-04 |
| Cer(d18:0_24:1) | 0.44 [0.16, 0.60] | 2.39E-15 [0.00, 0.00] | 0.56 [0.40, 0.76] | 0.44 [0.24, 0.60] | 0.22 [0.12, 0.33] | 0.009 | 1.00 | 2.68E-04 |
| Cer(d18:1_23:0) | 0.49 [0.00, 0.64] | 1.34E-13 [0.00, 0.38] | 0.51 [0.36, 0.70] | 0.49 [0.30, 0.64] | 0.25 [0.15, 0.43] | 0.049 | 1.00 | 1.95E-05 |
| Cer(d18:1_24:1) | 0.55 [0.20, 0.68] | 5.79E-15 [0.00, 0.28] | 0.45 [0.32, 0.63] | 0.55 [0.37, 0.68] | 0.27 [0.19, 0.41] | 0.006 | 1.00 | 8.41E-07 |
| Cer(d18:1_25:1) | 0.55 [0.07, 0.68] | 1.72E-14 [0.00, 0.38] | 0.45 [0.32, 0.63] | 0.55 [0.37, 0.68] | 0.27 [0.19, 0.46] | 0.025 | 1.00 | 6.58E-07 |
| Cer(d18:2_24:1) | 0.53 [0.17, 0.67] | 9.47E-16 [0.00, 0.28] | 0.47 [0.33, 0.65] | 0.53 [0.35, 0.67] | 0.27 [0.18, 0.41] | 0.009 | 1.00 | 2.97E-06 |
| DG(18:0_18:1) | 0.54 [0.10, 0.67] | 1.34E-14 [0.00, 0.38] | 0.46 [0.33, 0.62] | 0.54 [0.38, 0.67] | 0.27 [0.19, 0.46] | 0.019 | 1.00 | 1.15E-07 |
| DG(18:1_18:1) | 0.46 [0.04, 0.61] | 1.37E-14 [0.00, 0.35] | 0.54 [0.39, 0.72] | 0.46 [0.28, 0.61] | 0.23 [0.14, 0.40] | 0.034 | 1.00 | 1.74E-05 |
| DG(18:1_18:2) | 0.42 [0.12, 0.58] | 1.21E-14 [0.00, 0.23] | 0.58 [0.42, 0.77] | 0.42 [0.23, 0.58] | 0.21 [0.12, 0.34] | 0.014 | 1.00 | 1.76E-04 |
| PC(37:3) | 0.41 [0.13, 0.58] | 6.30E-15 [0.00, 0.20] | 0.59 [0.42, 0.78] | 0.41 [0.22, 0.58] | 0.21 [0.11, 0.32] | 0.012 | 1.00 | 4.21E-04 |
| PC(42:4e) | 0.37 [0.05, 0.55] | 1.65E-17 [0.00, 0.22] | 0.63 [0.45, 0.84] | 0.37 [0.16, 0.55] | 0.18 [0.08, 0.30] | 0.030 | 1.00 | 3.70E-03 |
| PE(18:0p_22:6) | 0.33 [0.00, 0.51] | 2.02E-14 [0.00, 0.23] | 0.67 [0.49, 0.88] | 0.33 [0.12, 0.51] | 0.16 [0.06, 0.29] | 0.048 | 1.00 | 1.16E-02 |
| TG(15:0_18:1_22:6) | 0.39 [0.10, 0.55] | 9.94E-17 [0.00, 0.22] | 0.61 [0.45, 0.80] | 0.39 [0.20, 0.55] | 0.20 [0.10, 0.31] | 0.016 | 1.00 | 5.00E-04 |
| TG(16:0_16:0_24:0) | 0.39 [0.12, 0.56] | 1.18E-16 [0.00, 0.19] | 0.61 [0.44, 0.81] | 0.39 [0.19, 0.56] | 0.20 [0.10, 0.31] | 0.012 | 1.00 | 8.63E-04 |
| TG(16:0_16:0_24:1) | 0.44 [0.03, 0.60] | 3.07E-13 [0.00, 0.33] | 0.56 [0.40, 0.74] | 0.44 [0.26, 0.60] | 0.22 [0.13, 0.38] | 0.039 | 1.00 | 9.41E-05 |
| TG(16:0_18:1_23:1) | 0.35 [0.00, 0.51] | 3.34E-14 [0.00, 0.27] | 0.65 [0.49, 0.84] | 0.35 [0.16, 0.51] | 0.17 [0.08, 0.31] | 0.048 | 1.00 | 2.42E-03 |
| TG(18:0_17:0_18:0) | 0.29 [0.01, 0.47] | 1.15E-14 [0.00, 0.20] | 0.71 [0.53, 0.91] | 0.29 [0.09, 0.47] | 0.14 [0.04, 0.26] | 0.044 | 1.00 | 2.03E-02 |
| TG(18:0_18:0_18:0) | 0.43 [0.14, 0.59] | 8.76E-15 [0.00, 0.21] | 0.57 [0.41, 0.77] | 0.43 [0.23, 0.59] | 0.21 [0.12, 0.33] | 0.010 | 1.00 | 1.95E-04 |
| TG(18:1_18:1_23:1) | 0.42 [0.10, 0.57] | 4.31E-15 [0.00, 0.26] | 0.58 [0.43, 0.77] | 0.42 [0.23, 0.57] | 0.21 [0.12, 0.34] | 0.019 | 1.00 | 1.55E-04 |
| TG(18:1_18:1_24:1) | 0.37 [0.09, 0.54] | 1.02E-15 [0.00, 1.00] | 0.63 [0.46, 0.83] | 0.37 [0.17, 0.54] | 0.19 [0.09, 0.30] | 0.019 | 1.00 | 1.76E-03 |
| TG(18:2_17:1_18:2) | 0.40 [0.16, 0.57] | 1.54E-15 [0.00, 0.17] | 0.60 [0.43, 0.79] | 0.40 [0.21, 0.57] | 0.20 [0.10, 0.30] | 0.007 | 1.00 | 4.35E-04 |
| TG(19:1_18:1_18:1) | 0.49 [0.17, 0.64] | 1.29E-14 [0.00, 0.26] | 0.51 [0.36, 0.68] | 0.49 [0.32, 0.64] | 0.25 [0.16, 0.38] | 0.009 | 1.00 | 4.53E-06 |

| | | | | | | | | |
|---|---|---|---|---|---|---|---|---|
| TG(19:1_18:1_18:2) | 0.40 [0.07, 0.56] | 2.48E-15 [0.00, 0.25] | 0.60 [0.44, 0.80] | 0.40 [0.20, 0.56] | 0.20 [0.10, 0.33] | 0.026 | 1.00 | 5.68E-04 |
| TG(25:0_16:0_18:1) | 0.45 [0.07, 0.60] | 4.17E-16 [0.00, 0.30] | 0.55 [0.40, 0.73] | 0.45 [0.27, 0.60] | 0.23 [0.13, 0.38] | 0.025 | 1.00 | 5.31E-05 |

Standardized additive genetic ($h^2$=heritability), shared environment ($h^2_C$) and unique environment ($h^2_E$) variance components (95% CI) of lipids were obtained using the ACE model. The columns p-AE, p-CE, and p-E, respectively, denote the p-values from the likelihood ratio test comparing ACE model vs AE, CE, and E models. p-CE is also the p value for heritability because testing the component A=0 is equivalent to testing that the heritability is zero. C.I. indicates confidence interval; DZ, dizygotic; ICC, intraclass correlation coefficient; MZ, monozygotic.

**Table S2: Heritability of summed lipid groups**

| Trait | A (95% C.I.) | C (95% C.I.) | E (95% C.I.) | ICC MZ (95% C.I.) | ICC DZ (95% C.I.) | p-CE | p-AE | p-E |
|---|---|---|---|---|---|---|---|---|
| Cer | 0.56 (0.19, 0.69) | 5.58E-14 (0.00, 0.29) | 0.44 (0.31, 0.61) | 0.56 (0.39, 0.69) | 0.28 (0.19, 0.43) | 0.007 | 1.00 | 5.56E-07 |
| Cer(d18:1/X) | 0.56 (0.16, 0.69) | 3.63E-14 (0.00, 0.31) | 0.44 (0.31, 0.62) | 0.56 (0.38, 0.69) | 0.28 (0.19, 0.43) | 0.010 | 1.00 | 6.34E-07 |
| Monounsaturated SM | 0.51 (0.07, 0.65) | 4.99E-15 (0.00, 0.35) | 0.49 (0.35, 0.67) | 0.51 (0.33, 0.65) | 0.26 (0.17, 0.42) | 0.027 | 1.00 | 6.36E-06 |
| Triglyceride | 0.48 (0.18, 0.62) | 1.07E-15 (0.00, 0.24) | 0.52 (0.38, 0.70) | 0.48 (0.30, 0.62) | 0.24 (0.15, 0.36) | 0.007 | 1.00 | 8.58E-06 |
| Polyunsaturated TG | 0.48 (0.03, 0.62) | 9.12E-14 (0.00, 0.37) | 0.52 (0.38, 0.69) | 0.48 (0.31, 0.62) | 0.24 (0.15, 0.42) | 0.036 | 1.00 | 5.40E-06 |
| PE | 0.46 (0.10, 0.62) | 9.96E-16 (0.00, 0.28) | 0.54 (0.38, 0.73) | 0.46 (0.27, 0.62) | 0.23 (0.14, 0.37) | 0.020 | 1.00 | 6.81E-05 |
| Cholesterol | 0.43 (0.07, 0.59) | 3.59E-15 (0.00, 0.26) | 0.57 (0.41, 0.78) | 0.43 (0.22, 0.59) | 0.21 (0.11, 0.34) | 0.025 | 1.00 | 4.97E-04 |
| HDL-C | 0.42 (0.03, 0.77) | 2.70E-01 (0.00, 0.61) | 0.31 (0.22, 0.44) | 0.69 (0.56, 0.78) | 0.48 (0.30, 0.65) | 0.035 | 0.24 | 8.00E-15 |
| CL_TG62 | 0.42 (0.05, 0.57) | 1.80E-14 (0.00, 0.30) | 0.58 (0.43, 0.77) | 0.42 (0.23, 0.57) | 0.21 (0.12, 0.36) | 0.032 | 1.00 | 1.60E-04 |
| LDL-C | 0.40 (0.12, 0.57) | 2.00E-17 (0.00, 0.20) | 0.60 (0.43, 0.80) | 0.40 (0.20, 0.57) | 0.20 (0.10, 0.31) | 0.013 | 1.00 | 8.73E-04 |
| CL_TG49 | 0.39 (0.00, 0.55) | 2.22E-16 (0.00, 0.32) | 0.61 (0.45, 0.80) | 0.39 (0.20, 0.55) | 0.19 (0.10, 0.36) | 0.048 | 1.00 | 3.81E-04 |

Standardized additive genetic (A=heritability), shared environment (C) and unique environment (E) variance components (95% CI) of lipids were obtained using the ACE model. The columns p-AE, p-CE, and p-E, respectively, denote the p-values from the likelihood ratio test comparing ACE model vs AE, CE, and E models. p-CE is also the p value for heritability because testing the component A=0 is equivalent to testing that the heritability is zero. C.I. indicates confidence interval; DZ, dizygotic; ICC, intraclass correlation coefficient; MZ, monozygotic. CL_TG49 and CL_TG62 represent sum of triglycerides with 44-49 total carbons, and 56-62 total carbons respectively while Cer(d18:1/X) represents sum of all ceramides with an 18:1 acyl chain in the sn-1 position .

**Figure S1: Genotypic correlation heatmap**

| | Cer | SM | TG | DG | PE | PC | LDL (Traditional lipids) | Chol (Traditional lipids) | HDL (Traditional lipids) | TG_t (Traditional lipids) |
|---|---|---|---|---|---|---|---|---|---|---|
| Cer | 0.94 | 0.83 | 0.34 | 0.32 | 0.38 | 0.245 | 0.43 | 0.45 | 0.09 | 0.36 |
| SM | 0.83 | N/A | 0.39 | 0.17 | 0.42 | 0.16 | 0.4 | 0.41 | 0.183 | 0.189 |
| TG | 0.34 | 0.19 | 0.81 | 0.7 | 0.4 | 0.39 | 0.38 | 0.34 | -0.28 | 0.72 |
| DG | 0.32 | 0.17 | 0.7 | 0.73 | 0.33 | 0.33 | 0.54 | 0.45 | -0.37 | 0.8 |
| PE | 0.38 | 0.42 | 0.4 | 0.33 | 0.86 | 0.41 | 0.4 | 0.43 | 0.43 | 0.06 |
| PC | 0.25 | 0.16 | 0.39 | 0.33 | 0.41 | 0.41 | 0.4 | 0.41 | 0.36 | 0.48 |
| LDL (Traditional lipids) | 0.43 | 0.4 | 0.38 | 0.54 | 0.4 | 0.4 | | | | |
| Chol (Traditional lipids) | 0.45 | 0.41 | 0.34 | 0.45 | 0.43 | 0.41 | | | | |
| HDL (Traditional lipids) | 0.09 | 0.183 | -0.28 | -0.37 | 0.43 | 0.36 | | | | |
| TG_t (Traditional lipids) | 0.36 | 0.189 | 0.72 | 0.8 | 0.06 | 0.48 | | | | |

Genetic correlation matrix heatmap. Values represent the median of genetic correlations taken between combinations of heritable lipid species of one lipid class with lipid species of another class (or the same class). Note SM represents the sum of SM with a single double bond, thus no correlation could be computed for SM with itself. TG_t represents triglycerides referred to as a traditional lipid measure (as opposed to individual species measured by mass spectrometry).

**Figure S2**

**A)**

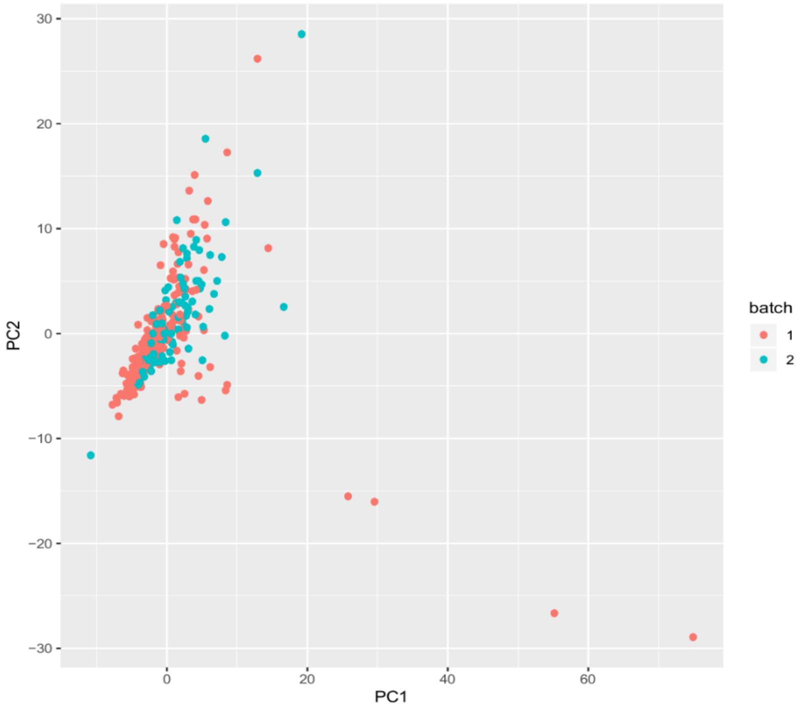


B)

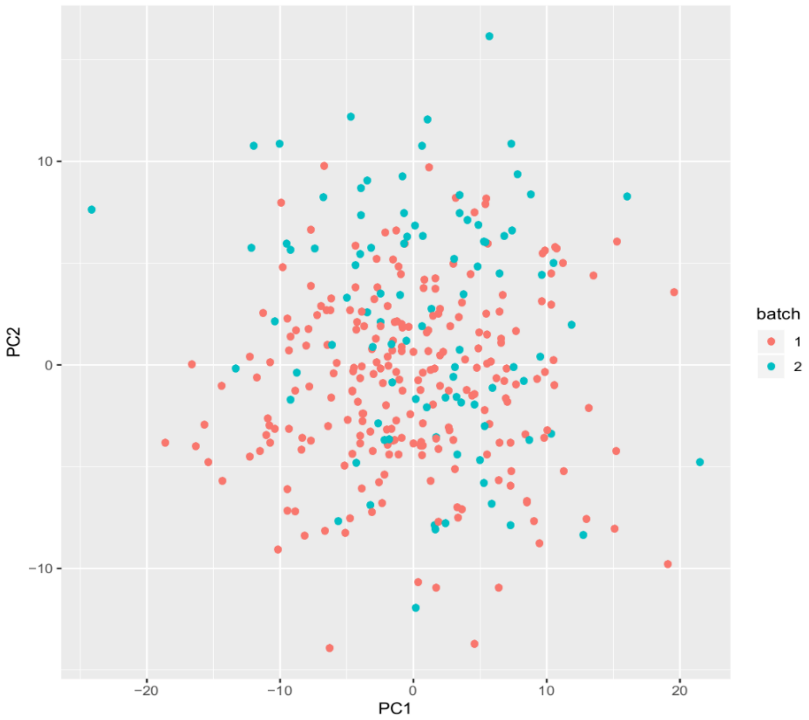


PCA plots showing good overlap of experimental batch lipids after A) residuals were taken and B) after inverse rank normal transformation was applied to these residuals.

**Table S3**

Lipid (triglyceride)-gene expression associations listed by heritability and degree of saturation

| | Heritable | Non heritable |
|---|---|---|
| Saturated TG | *HDC*<br>*ADAM8, APMAP, CPA3, RP11-179G5.1, RPL4P2* | *APMAP, CPA3, HDC, MGAM*<br>*GATA2, KRT23* |
| Mono-unsaturated TG | *CPA3, LILRA6, LILRB3, SNHG1, TRIM51*<br>*GAB2, HDC, PRKCD, RP11-350G8.3, RPSA, RPSAP15, RPSAP58, SNORA62* | *HDC*<br>*CPA3, GATA2, SLC45A3* |
| Polyunsaturated TG | *HDC*<br>*VAMP8*<br>*CPA3, GAB2, LILRA6, LILRB3, PRKCD, REPS2, RP11-179G5.1, RPL35P5, RPS10P14* | *HDC*<br>*CPA3*<br>*SLC45A3* |

This table includes gene transcripts associated with ≥3 lipids in each saturation class, and also includes the transcripts associated with the third highest number of lipids among the polyunsaturated TG class.

Abbreviations of gene names are based on Gene Ontology nomenclature:

ADAM8, A disintegrin and metalloproteinase domain-containing protein 8; APMAP, Adipocyte plasma membrane-associated protein; CPA3, carboxypeptidase A3; FPR1, Formyl peptide receptor 1; GAB2, GRB2-associated-binding protein 2; GATA2, Endothelial transcription factor GATA-2; HDC, Histidine decarboxylase; KRT23; Keratin, type I cytoskeletal 23; LILRA6, Leukocyte immunoglobulin-like receptor subfamily A member 6; LILRB3Leukocyte immunoglobulin-like receptor subfamily B member 3; MGAM, Maltase-glucoamylase; PRKCD, Protein kinase C delta type; REPS2, RalBP1-associated Eps domain-containing protein 2; RP11.179G5.1, Ribosomal Protein SA Pseudogene 18; RP11.350G8.3, Ribosomal Protein SA Pseudogene 17; RPL35P5, Ribosomal Protein L35 Pseudogene 5; RPL4P2, Ribosomal Protein L4 Pseudogene 2; RPS10P14, Ribosomal Protein S10 Pseudogene 14; RPSA, 40S ribosomal protein SA; RPSAP15, Ribosomal Protein SA Pseudogene 15; RPSAP58, Ribosomal protein SA pseudogene 58; S100A11P1, S100 Calcium Binding Protein A11 Pseudogene 1; SLC45A3, Solute carrier family 45 member 3; SNHG1, Small Nucleolar RNA Host Gene 1; SNORA62, 40S ribosomal protein SA; TRIM51, Tripartite motif-containing 51; VAMP8, Vesicle-associated membrane protein 8.

#### Supplemental data

Supplemental Data S1-S8 is available as sheets from an Excel file accessable through this link:

https://unsw-my.sharepoint.com/:x:/g/personal/z3375126_ad_unsw_edu_au/EV_Io0TjE3VDseCzE6KwFT0BIWRCpUB5DBAMZeBhK-vfNQ?e=exlr1n
